# Cerebral hypoperfusion exacerbates traumatic brain injury in male but not female mice

**DOI:** 10.1101/2023.10.19.563077

**Authors:** Bailey J. Whitehead, Deborah Corbin, Megan L. Alexander, Jacob Bumgarner, Ning Zhang, A. Kate Karelina, Zachary M. Weil

**Affiliations:** Department of Neuroscience and Rockefeller Neuroscience Institute, West Virginia University, Morgantown WV USA

## Abstract

Mild-moderate traumatic brain injuries are common and while many individuals recover fully there is mounting clinical and epidemiological evidence that for a substantial subset, even when the acute TBI symptoms resolve, long term health can still be impacted. Individuals with a history of TBI are disproportionately vulnerable to many disease conditions including age-related neurodegeneration. These relationships are difficult to predict but these outcomes very likely interact with other disease risk factors such as cardiovascular disease. Here we tested the hypothesis that a mild pre-injury reduction in cerebral blood flow (bilateral carotid artery stenosis; BCAS) would impair recovery from TBI. Male and female mice underwent BCAS using steel microcoils around the carotid arteries, a mild-moderate closed-head TBI, or a combination of BCAS followed by TBI 30 days post-implantation. Cerebral blood flow, spatial learning and memory, axonal damage, and gene expression profiles were assessed. BCAS led to a ∼10% reduction in CBF, while TBI caused a similar decrease. However, mice exposed to both BCAS and TBI exhibited more pronounced reductions in CBF, associated with marked spatial learning and memory deficits, particularly in males. Axonal damage in male mice was also exacerbated by the combination of BCAS and TBI compared to either injury alone. Females exhibited spatial memory deficits associated with BCAS, but this was not exacerbated by TBI. We performed single nuclei RNA sequencing on male brain tissue to investigate the mechanisms underlying poorer long term functional outcomes in in TBI-BCAS animals. TBI and BCAS independently altered gene expression profiles in neurons and glia but in most cases BCAS and TBI together produced markedly different transcriptional patterns than either challenge alone. Overall, our findings reveal that the presence of mild reductions in cerebrovascular blood flow as a proxy for preexisting cardiovascular disease significantly exacerbated TBI outcomes in male but not female mice, indicating that even relatively mild comorbidities could significantly alter TBI outcomes and increase the probability of secondary disease processes.

## Introduction

Classically, mild-to-moderate traumatic brain injuries (TBI) and the associated recovery were considered to be isolated events and were believed to rarely yield long term sequela (1). However, recent clinical and epidemiological data have strongly indicated that, even when the acute TBI symptoms resolve, chronic medical challenges can endure (2, 3). Indeed, individuals with a history of brain injury are more susceptible to certain pathologies including epilepsy, depression, renal dysfunction, cardiometabolic disease, stroke, and neurodegeneration in the years after their injury; in addition, morbidity and mortality from all causes is higher among these individuals as they age (4).

TBI is a major risk factor for the development of age-related neurodegeneration. Indeed, systematic reviews of TBI across the severity spectra have reported a ∼60-90% elevation in all cause dementia among former TBI patients (4, 5). The causes of this phenomenon remain incompletely understood and challenging to predict. TBI is known to interact with other risk factors to further exacerbate disease vulnerability. For instance, among US veterans, dementia risk is increased in individuals with cardiovascular disease and individuals with a history of TBI. However, the combination of brain injury in individuals with ongoing cardiovascular disease greatly increased the overall risk (4). Since cardiovascular disease, metabolic dysfunction, and associated risk factors such as smoking and hypertension are all common among TBI survivors, there is an urgent need to understand how these variables interact to alter neurodegenerative outcomes (6–8)

Moreover, it is becoming increasingly clear that preexisting disease states can alter both the incidence and consequences of TBI. For instance, both cerebrovascular disease and depression predicted the incidence of TBI in older adults and vascular dysfunction increased the likelihood that injury resulted in death (9). Similarly, preinjury hyperlipidemia strongly predicted the development of new onset anxiety disorders after TBI (10). Indeed, it has been argued that the vast majority of TBI patients (more than 97%) have some form of comorbidity that both potentially alters disease outcomes and limits their inclusion in many clinical studies (11).

Here we focus on the strong relationship among cerebral blood flow, traumatic brain injuries and neurodegeneration. The nervous system has several features that render it disproportionally vulnerable to disruption in blood flow. Nervous tissue is energetically expensive to maintain, and energy costs rise with synaptic activity, yet neurons and glia store very little energy (12). Thus, the cerebral vasculature must alter the delivery of oxygen and glucose (among other substrates and waste removal functions) in concert with neuronal activity. Disease states that impair cerebral blood flow or the activity dependent coupling of blood flow have the potential to significantly impair neuronal function and this cerebrovascular dysfunction has emerged as a key mechanism of neurodegenerative disease (13, 14). Moreover, a variety of risk factors for neurodegenerative diseases including age, smoking, obesity, diabetes, hypertension, and APOE4 genotypes are linked with a reduction in cerebral blood flow (15). Chronic reductions in cerebral blood flow are associated with decrements in cognitive function, constraints on neuroplasticity and impairments in central metabolic function in both humans and experimental animals (16–19).

It has long been known that a key component of the pathophysiology of TBI involves vascular dysfunction (20). TBI can result in direct mechanical damage to vessels and secondary injuries associated with inflammation and metabolic dysfunction in vascular cells. Additionally, individuals with a history of TBI very often experience autonomic dysfunction that can further imperil cardio- and cerebrovascular regulation and in general coupling of blood flow to neuronal activity is disrupted by injury(21). A majority of TBI patients exhibit overall reductions in cerebral blood flow in the hours after TBI resulting in a general disconnect between metabolic needs of the tissue and blood flow(22, 23). What is less well understood is how preexisting vascular dysfunction can modulate TBI outcome. Here we tested the hypothesis that mild but chronic reductions in cerebral blood flow (as a proxy for preexisting cardiovascular risk factors) prior to injury would exacerbate tissue damage, impair cerebral blood flow, and produce poorer functional outcomes in mice. Specifically, we injured adult mice of both sexes 30 days after implantation of arterial coils and then assessed their cerebral blood flow, spatial learning and memory performance, transcriptional activity at the single nuclei level, and inflammatory responses.

## Methods

All procedures were approved by the West Virginia University Institutional Animal Care and Use committee and were conducted in accordance with NIH guidelines. Swiss Webster mice derived from progenitors acquired from Charles River were bred in our colony at WVU. Mice of both sexes were randomly assigned to receive either the bilateral carotid artery stenosis (BCAS) procedure or the SHAM condition. Thirty days post BCAS or SHAM, groups were again divided in half and animals received either the TBI or control procedure (Fig 1). This produced four groups of animals per sex SHAM, TBI, BCAS, and TBI-BCAS.

**Fig 1.**
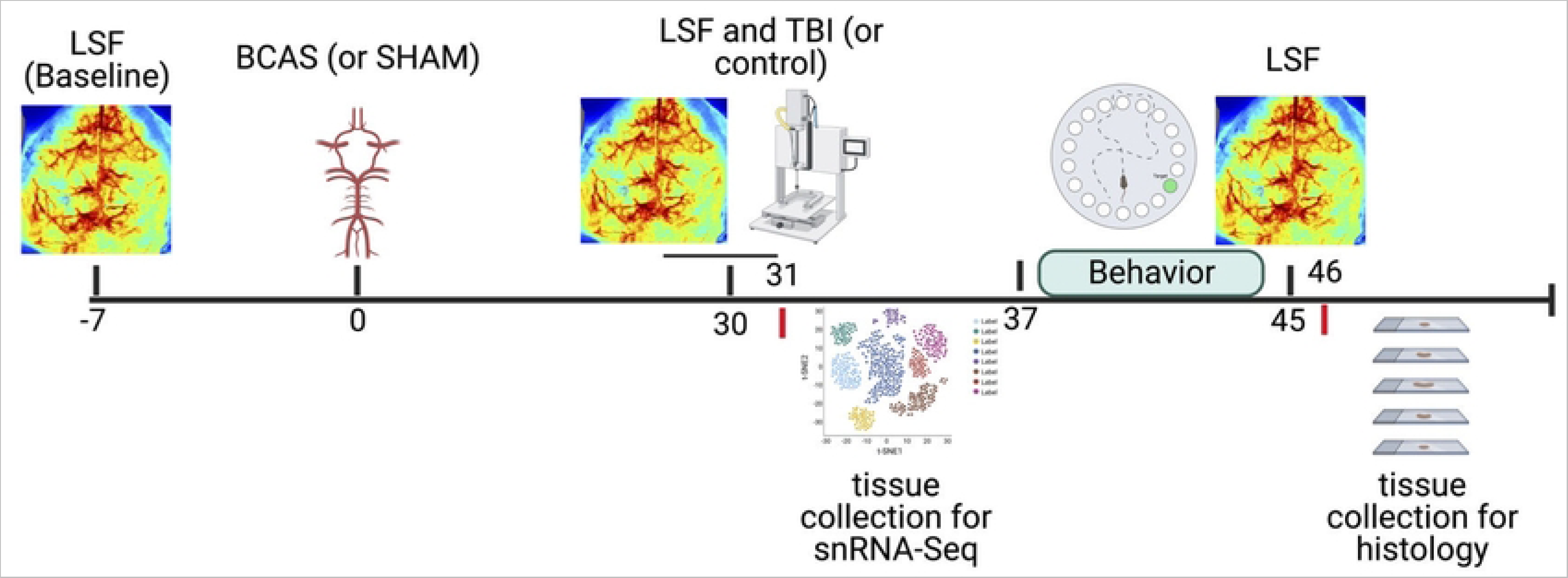
Timeline of experimental procedures. All mice underwent baseline cerebral blood flow (CBF) analysis followed by BCAS (or sham procedure) 7 days later. Thirty days later, mice underwent a second CBF analysis and were further split into two groups, receiving a TBI or sham procedure. In one cohort, mice underwent behavioral testing, followed by a final CBF analysis, and brain tissue was collected for histology. In another cohort, brain tissue was dissected 24 hours after TBI or sham injury for single nuclei RNA sequencing.

### BCAS Procedure

Swiss Webster mice of both sexes (approximately 10 weeks of age) were deeply anesthetized, local analgesia was provided (1.5 mg/kg bupivacaine and 0.5 mg/kg lidocaine), and a midline cervical incision made to expose the carotids. The arteries were gently freed from their sheaths and stainless steel (0.08 diameter wire; SWP-A) microcoils (internal diameter=0.2 mm, pitch=0.05 mm, total length=2.5 mm)(24) placed around the carotid arteries on both sides below the junction of the internal and external carotid artery branches. The control procedure was identical to the BCAS except that the coils were not implanted. The skin was sutured, and the animal allowed to recover in its home cage.

### Mild-Moderate TBI procedure

A single closed-head mild-moderate traumatic brain injury was conducted as previously described (25). Mice were randomly assigned to experimental group and then placed into a stereotaxic frame under 1.5% inhaled isoflurane anesthesia, local analgesia was provided (1.5 mg/kg bupivacaine and 0.5 mg/kg lidocaine), and an incision was made down the midline of the scalp. A 5mm diameter impactor tip was retracted and driven onto the skull along the midline (−2mm AP relative to bregma) to a depth of 1.2 mm at a rate of 5mm/sec (dwell time: 30 msec). The control procedure was performed identically except that the impactor was not driven into the skull (just placed lightly on the surface and then retracted). Mice were treated with 5.0 mg/kg meloxicam, the skin was sutured, and the animal was allowed to recover in its home cage.

### Laser Speckle imaging

Cerebral blood flow was acquired via a MoorFLPI Laser Doppler system. Briefly, mice were anesthetized with isoflurane, an incision was made to expose the skull, and 3 images acquired over 2 minutes. The incision was treated with a local anesthetic (1.5 mg/kg bupivacaine and 0.5 mg/kg lidocaine) and sutured. Subsequent measurements were performed by reopening the scalp at the same incision site and repeating the same procedure. Blood flow imaging took place three times over the course of the experiment, seven days prior to BCAS (baseline), 30 days post BCAS (immediately before TBI procedure), and again 15 days after the TBI. All values were normalized to the baseline.

### Barnes Maze

The Barnes maze is a brightly lit circular arena (36 inches in diameter) with 18 evenly spaced holes, and one leading to a dark box. Various shapes/colors positioned around the maze served as visual cues to the location of the escape box. Mice underwent five days of training consisting of three consecutive 120-sec trials separated by brief returns to their home cages. One day after the last training trial, animals were given a 90-sec probe trial in which the escape box was removed. Latency to find the target hole and time spent in the quadrant that formerly contained the target recorded during the probe trial were video recorded daily and analyzed using AnyMaze software.

### Single Nuclei RNA Sequencing

Males from each of the groups were anesthetized and decapitated 24 hours post TBI. An approximately 2mm cube of forebrain tissue was dissected under the TBI and frozen in liquid nitrogen. Frozen tissue was then submitted to Genewiz for 10X Genomics snRNA-SEQ on the 10X Genomics Chromium System. Single nuclei RNA libraries were generated using the Chromium Single Cell 3’ kit (10X Genomics, CA, USA). Loading onto the Chromium Controller was performed to target capture of ∼3,000 GEMs per sample for downstream analysis and processed through the Chromium Controller following the standard manufacturer’s specifications. The sequencing libraries were evaluated for quality on the Agilent TapeStation (Agilent Technologies, Palo Alto, CA, USA), and quantified using Qubit 2.0 Fluorometer (Invitrogen, Carlsbad, CA). Pooled libraries were quantified using qPCR (Applied Biosystems, Carlsbad, CA, USA) prior to sequencing using Illumina 2 x 150bp at a target death of 50,000 pair-end end reads per cell at a configuration compatible with the recommended guidelines as outlined by 10X Genomics. Cell Ranger was used to align reads to the Mus musculus genome build GRCm38 and produce a filtered gene x cell matrix of UMI counts. The filtered matrix was then used as input for the Seurat R software package to create a Seurat R object. Samples were then filtered to exclude cells that contained greater than 5% mitochondrial gene contamination (26). Gene counts were then normalized, and batch correction performed using the Harmony (27) algorithm. Nuclei were clustered using Louvain algorithm (0.1 and 0.3 resolution) and annotated with SCSA(28) using the 2023 CellMarker database. Doublets were removed with DoubletFinder (29). Gene set expression analysis was performed using cell clusters defined by 0.3 resolution clustering.

Seurat objects were then loaded into BbrowserX (Bioturing, San Diego, California) for downstream analysis. Differentially expressed genes (DEGs) were detected by comparing clusters between groups using the Venice algorithm (30). For all cell types four comparisons were conducted: SHAM V. TBI, SHAM V. BCAS, SHAM V. TBI-BCAS, and TBI V. TBI-BCAS. DEGs were genes that had false discovery rates (FDR) of less than 0.05 and differed by at least ± 0.25 Log fold change. This analysis produced eight lists of genes (up- and down-regulated genes in each comparison). These lists were used to produce a multi-query gene ontology search in Gprofiler and resulting ontologies were plotted(31). AUCell analysis was also conducted for each of the comparisons which uses Wikipathways gene sets as comparators (32).

### Histological Staining

Following the conclusion of the Barnes maze probe and final Laser Speckle Flowmetry session, animals were euthanized and transcardially perfused. Following perfusion, brain tissue was cryopreserved, frozen and sliced on a cryostat at 40µm throughout the forebrain. The sliced tissue was then used for free-floating silver staining (NeuroSilver, FD Neurotechnologies, Columbia, MD) per the manufacturer’s instructions.

### Imaging and Analysis

Axonal degeneration was assessed via silver staining by an investigator blinded to experimental conditions on an Olympus VS-120 microscope with a 10x Plan S Apo/0.4 NA objective using a Monochrome XM10 camera (1376 x 1032 imaging array, 6.45 x 6.45-µm pixel size, and 14-bit digitization). Experimenters made qualitative analyses of silver staining using a 4-point scale (0 = no silver staining, 3 = dense axonal degeneration) (33). Silver staining images were collected and analyzed in 4 regions of interest: corpus callosum, optic chiasm, thalamus, and internal capsule.

### Data Analysis

Effect sizes were >0.45 with biochemical and histological changes being larger than behavioral effects. Thus, we powered the study to detect differences with a β>0.8. Behavioral outcomes were assessed via a two-way ANOVA (blood flow x injury) and were conducted separately for both sexes. All significant overall results (p < 0.05) were followed up with an LSD post hoc analysis. Qualitative silver analyzed using the non-parametric Kruskal-Wallis H test.

## Results

### Blood Flow

To determine whether BCAS impaired cerebral blood flow we performed laser speckle flowmetry imaging and compared values to those taken prior to implanting restrictive coils around the carotids. In both males (F_1,29_=9.34, p<0.005; Fig 2), and females (F_1,33_=5.64, p<0.05), BCAS reduced cerebral blood flow. Next, we injured the animals with mild/moderate closed head injuries or the control procedure and then reassessed cerebral blood flow 15 days later. At the final time point both BCAS (F_1,29_=19.383, p<0.001), and TBI (F_1,29_=15.025, p<0.001) strongly reduced cerebral blood flow in males. This produced an additive effect such that BCAS+TBI animals differed significantly from all other groups (p<0.05 in all cases) though in no cases were the interactions significant. However, at this timepoint, the overall effect of BCAS was no longer significant among females ((F_1,29_=4.018, p=0.054), though TBI did reduce blood flow (F_1,29_=9.40, p<0.005). Moreover, while BCAS+TBI had the lowest numerical blood flow it did not differ significantly from TBI among the females (p=0.07).

**Fig 2.**
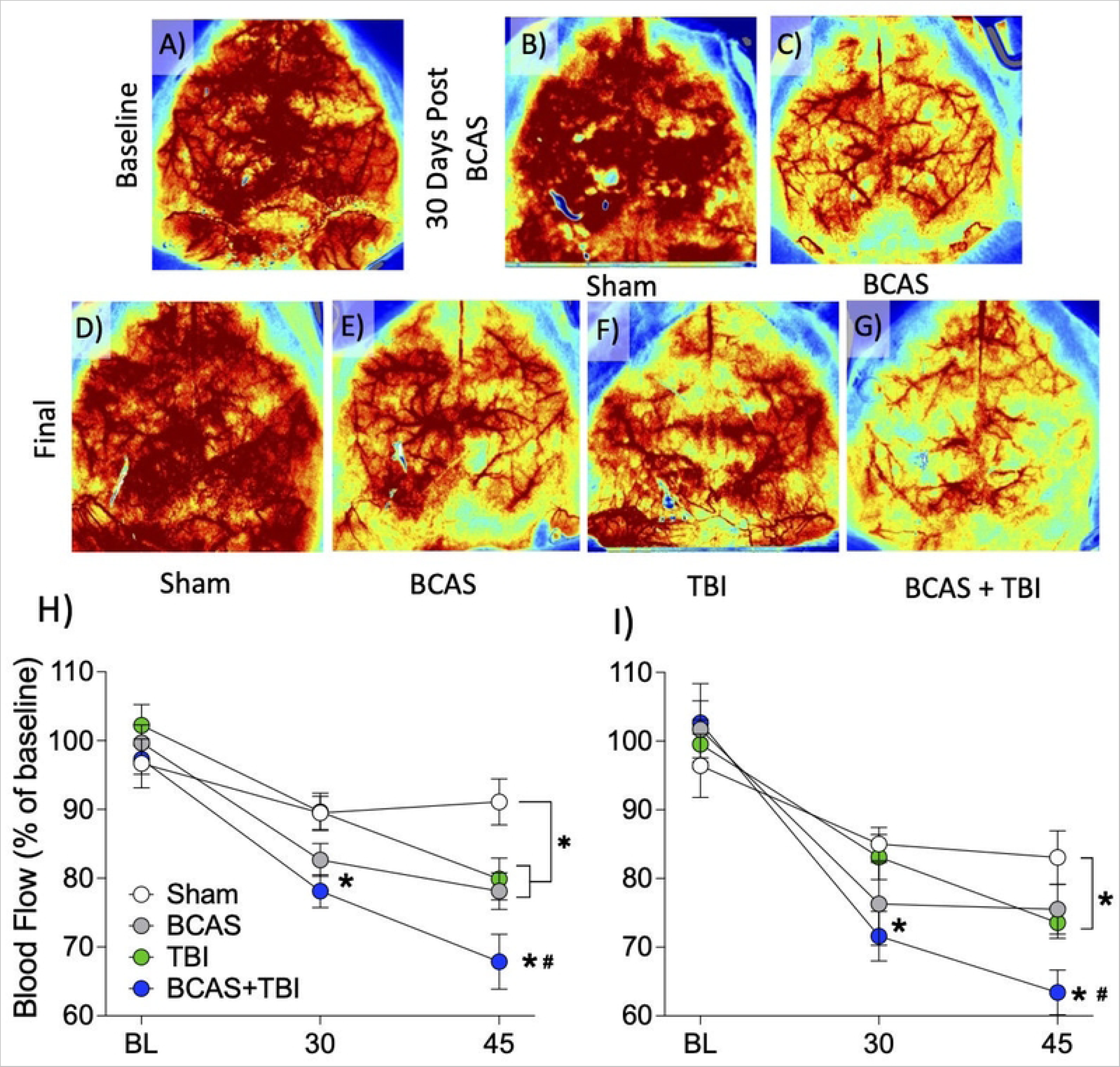
BCAS and TBI impair cerebral blood flow. Representative cerebral blood flow (CBF) is shown A) at baseline, B) 30 days following sham hypoperfusion, or C) 30 days following BCAS. Immediately after the 30-day CBF assessment mice underwent TBI or another sham surgery, and a final measure of CBF was reassessed 15 days later; D) control, E) BCAS only, F) TBI only, G) BCAS + TBI. CBF was quantified and reported as percent of baseline at each time point in H) male and I) female mice. BCAS produced a moderate reduction in cerebral blood flow at thirty days that was significantly exacerbated in mice that underwent a TBI. Notably TBI also produced persistent reduction in blood flow in the absence of carotid manipulation. Laser speckle flowmetry (warmer colors indicate greater blood flow) was used to assess cerebral blood flow prior to any surgical manipulations. N=8-10 mice group and * p<0.05 significantly different from sham, # indicated different from TBI. Data are presented as mean (±SEM).

### Spatial Learning and Memory

The effects of TBI on spatial learning and memory were modulated by BCAS. A repeated measures analysis revealed a between-subjects effect on latency to escape in the Barnes maze in male (F_3, 31_ = 3.647, p < 0.05; Fig 3) but not female mice (F_3,29_ = 1.675, p = 0.194). A post hoc analysis in male mice revealed that while neither TBI nor BCAS alone impaired learning and memory, the combination of BCAS+TBI significantly impaired performance on the Barnes maze relative to control and BCAS alone (p < 0.05). Among females, repeated measures analysis revealed a simple main effect of BCAS (F_1,29_ = 4.89, p < 0.05) on learning and memory but not TBI (F_1,29_ = 0.00, p = 1.00), nor an interaction of BCAS+TBI (F_1,29_ = 0.235, p = 0.632). Moreover, among males, TBI reduced the time spent in the target quadrant during the probe trial (F_1,31_= 6.307, p<0.05) but there was no significant effect of BCAS (F_1,31_=1.087, p>0.05) and the interaction between the variables was not significant. In contrast, among females, there were no significant effects of either TBI (F_1,29_=0.115, p>0.05) or BCAS (F_1,29_=3.883, p>0.05) in the probe trial.

**Fig 3.**
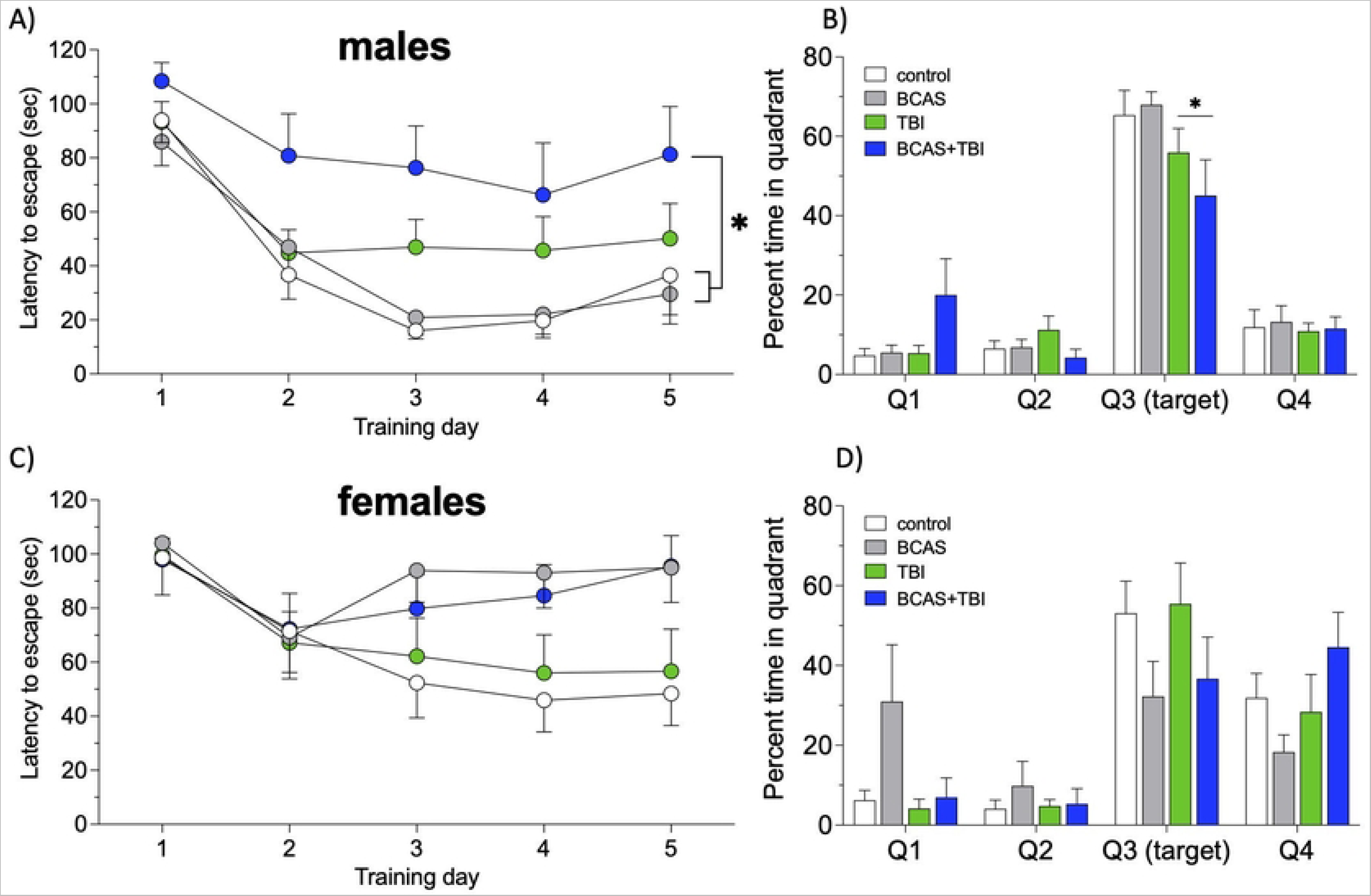
BCAS exacerbates TBI-induced cognitive dysfunction in males. Barnes maze performance was slightly impaired by TBI (numerical but not statistical increase in escape latency A) and reduction in time in escape quadrant during probe trial B). However, TBI-BCAS mice exhibited significantly worse cognitive performance. C-D) among female BCAS impaired Barnes maze performance but was not altered by injury. N=8-10 mice/group. Testing was initiated one week following TBI procedure. * Indicates significantly different from control animals.

### Histopathology

Traumatic brain injury damaged axons across the forebrain as assessed by qualitative analysis of silver staining. Specifically, among males, there was an overall surgery effect (U_3,37_=18.881, p<0.0001) which was mediated by greater degeneration among TBI groups. Multiple comparison testing revealed that BCAS+TBI mice had greater axonal degeneration compared to TBI alone (p<0.05; Fig 4). In contrast, among females only TBI altered silver staining (U_3,40_=18.435, p<0.0001) but there was no difference between TBI and BCAS+TBI (p>0.05).

**Fig 4.**
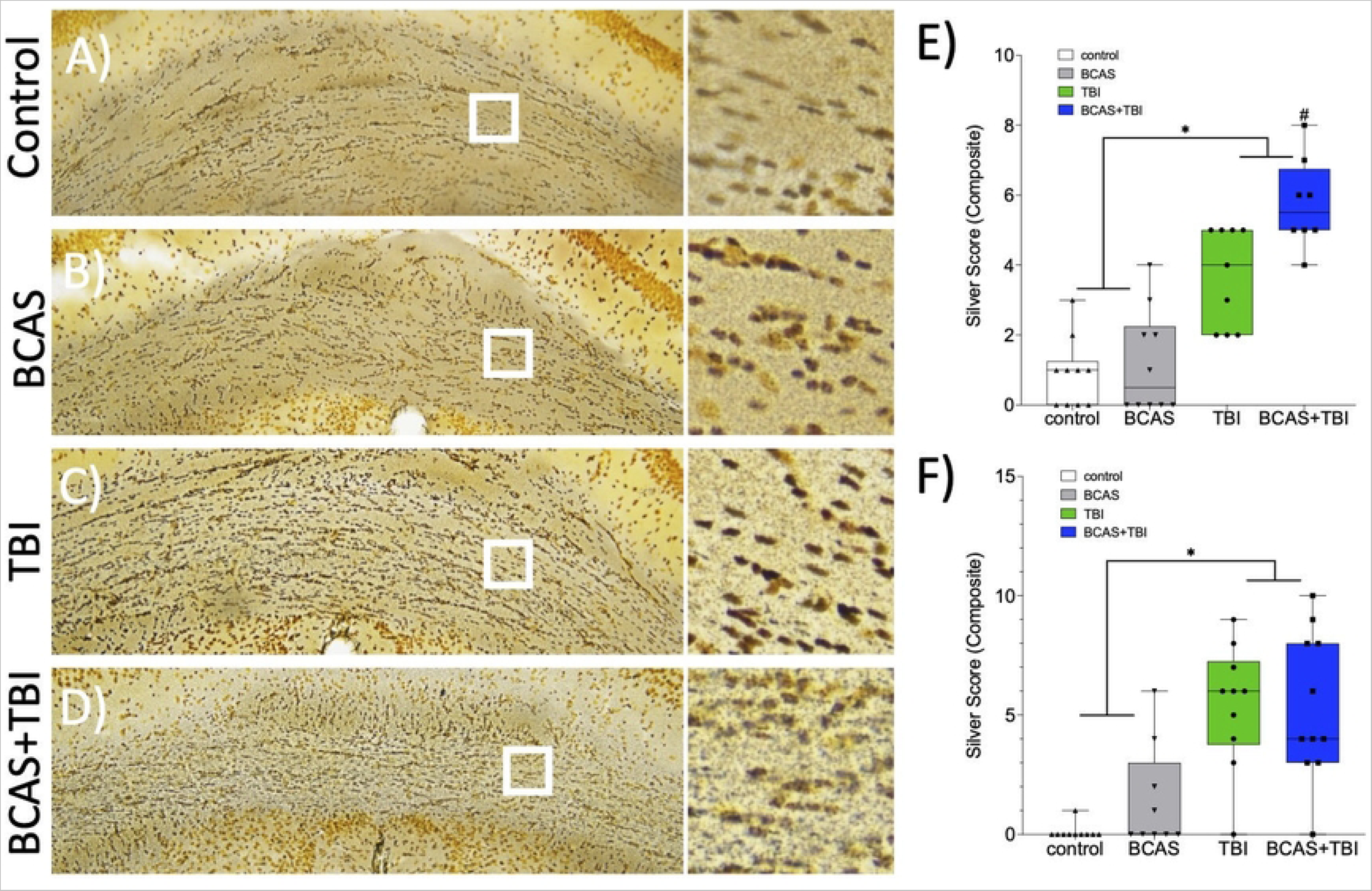
BCAS exacerbates axonal damage following TBI in males. Silver staining in the corpus callosum of A) control, B) BCAS, C) TBI, and D) BCAS-TBI mice. White boxes indicate insets. BCAS exacerbates axonal degeneration among males E) but not females). * Indicates significantly different from control animals, # indicates different from TBI.

### RNA Sequencing

Louvain (0.3) algorithm-based clustering produced 13 clusters of cells (astrocyte, CCK basket cell, cortical layer cells, endothelial cells, ependymal cells, GABAergic neurons, glutamatergic neurons, medium spiny neurons, microglia, neurons, oligodendrocytes, and smooth muscle cells). Neuron clusters were manually recombined based on expression of markers for glutamatergic (SLC17A6, SLC17A7) and GABAergic activity (SLC32A1, GAD2). Cluster defining genes are shown in Fig 5 and included in supplementary data.

**Fig 5.**
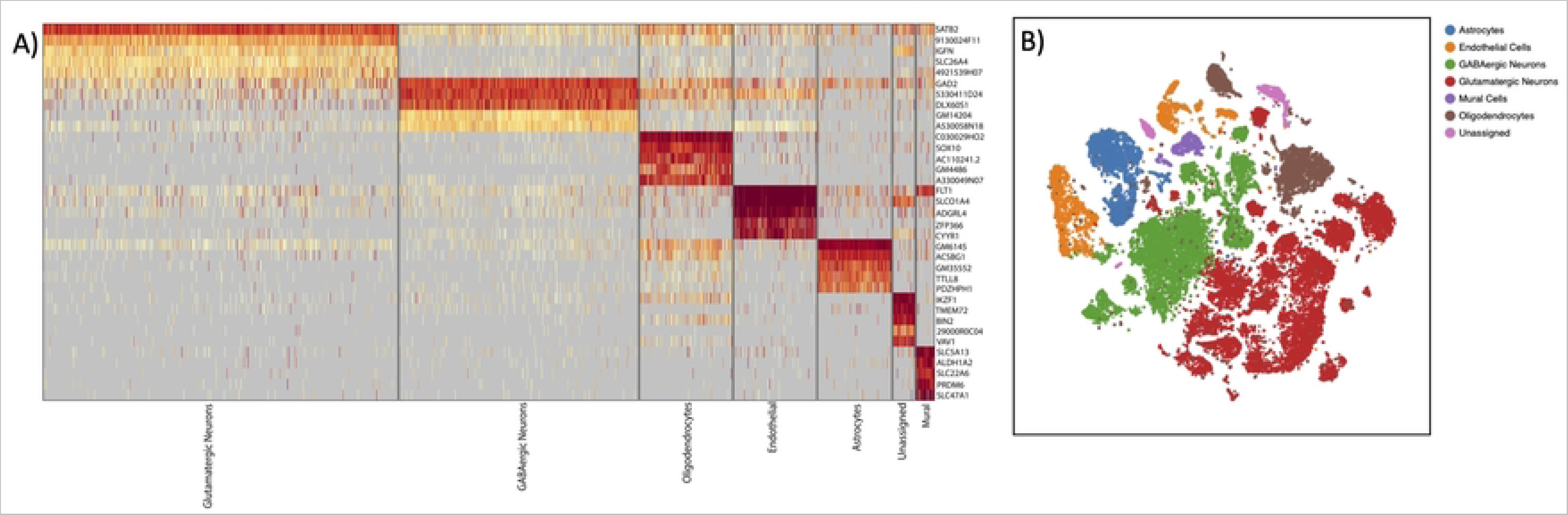
Single nuclei RNA sequencing. A) heatmap of cluster-defining genes, B) TSNE plot of clustered cells.

#### Glutamatergic Neurons

BCAS significantly altered transcriptional activity both in TBI and SHAM-injured animals. Specifically, BCAS alone upregulated genes were largely related to intracellular signal transduction and developmental processes (Fig 6). Notably, there were over 250 genes significantly downregulated by BCAS. These genes encoded proteins responsible for growth factor signaling, control of RNA transcription, and general cellular biosynthetic activity. Remarkably there was a relatively minor transcriptional response to TBI alone in these cells.

**Fig 6.**
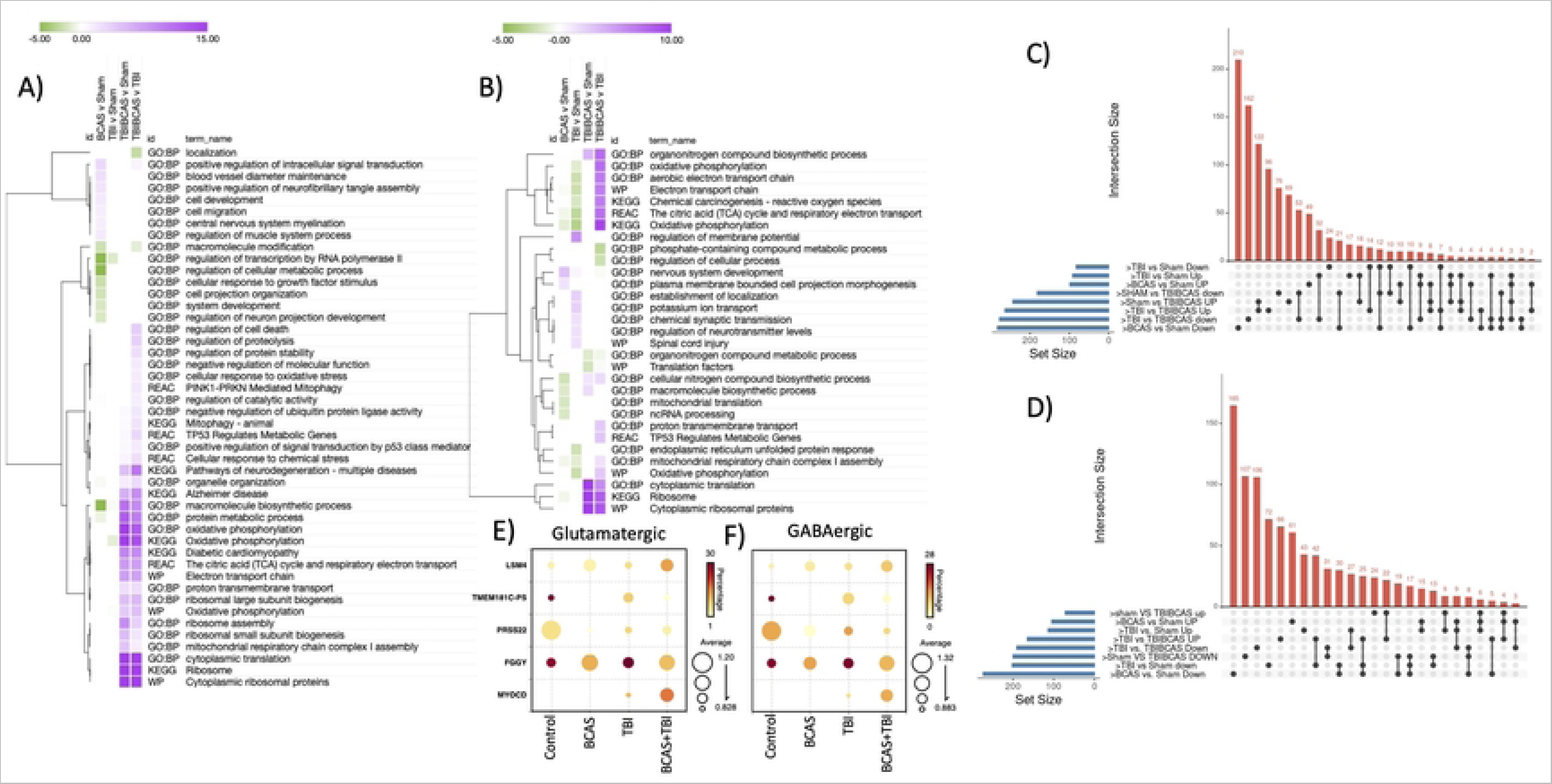
BCAS and TBI alter gene expression in neurons. Heatmap of gene ontology terms derived from differentially expressed genes in A) glutamatergic and B) GABAergic neurons. Upset plots describing the relationships between gene lists in C) glutamatergic and D) GABAergic neurons and notable genes altered by treatment in E-F).

Gene ontology analysis revealed a small reduction in biosynthesis-related genes. This is in sharp contrast to the transcriptional response among BCAS+TBI animals compared to both SHAM and TBI alone. This group exhibited dramatic upregulation of genes related to oxidative phosphorylation, ribosome subunit metabolism, P53 mediated stress signaling, as well as genes related to proteolysis, response to cellular stress and programmed cell death as well as the KEGG term associated with Alzheimer’s Disease.

#### GABAergic Neurons

BCAS also significantly altered transcriptional responses in GABA neurons. In a manner that was generally similar to excitatory neurons, BCAS downregulated GABA gene expression associated with large molecular biosynthesis. However, BCAS also downregulated the expression of genes associated with oxidative metabolism, mitochondrial proteins, and ribosome subunit expression. TBI alone also downregulated mitochondria and electron transport chain -related genes, as well an upregulated the expression of genes associated with maintaining membrane potentials. Once again, the combination of BCAS+TBI resulted in DEGs that differed markedly from either condition alone. Again, many of the genes upregulated in this group were related to ribosome production and components of the electron transport chain. Remarkably there were several gene ontologies where TBI and BCAS+TBI produced opposite transcriptional effects. Thus, the comparison between TBI and BCAS+TBI often produced larger effects (e.g., on oxidative phosphorylation) then the comparison between BCAS+TBI and SHAM because TBI downregulated oxidative phosphorylation-related genes (relative to SHAM) unless mice had also experienced BCAS, in which case these genes were strongly upregulated.

#### Oligodendrocytes

Oligodendrocyte responses to BCAS and TBI differed significantly from neurons. BCAS-mediated hypoperfusion significantly upregulated genes related to cellular localization, synapse formation, learning and memory, response to neurotrophins, and cell adhesion molecules among many others (Fig 7). Myelination related genes were reduced by BCAS. TBI, in contrast, upregulated genes associated with cellular migration, cell adhesion, synaptic signaling, and ribosome production. Many of these oligodendrocyte transcriptional changes that involve plasticity and support for neurons and synapses are not present in cells from BCAS+TBI animals. The largest numbers of downregulated genes are in the comparison between TBI and BCAS+TBI animals where many genes associated with transcriptional activity and ribosome biogenesis are significantly inhibited.

**Fig 7.**
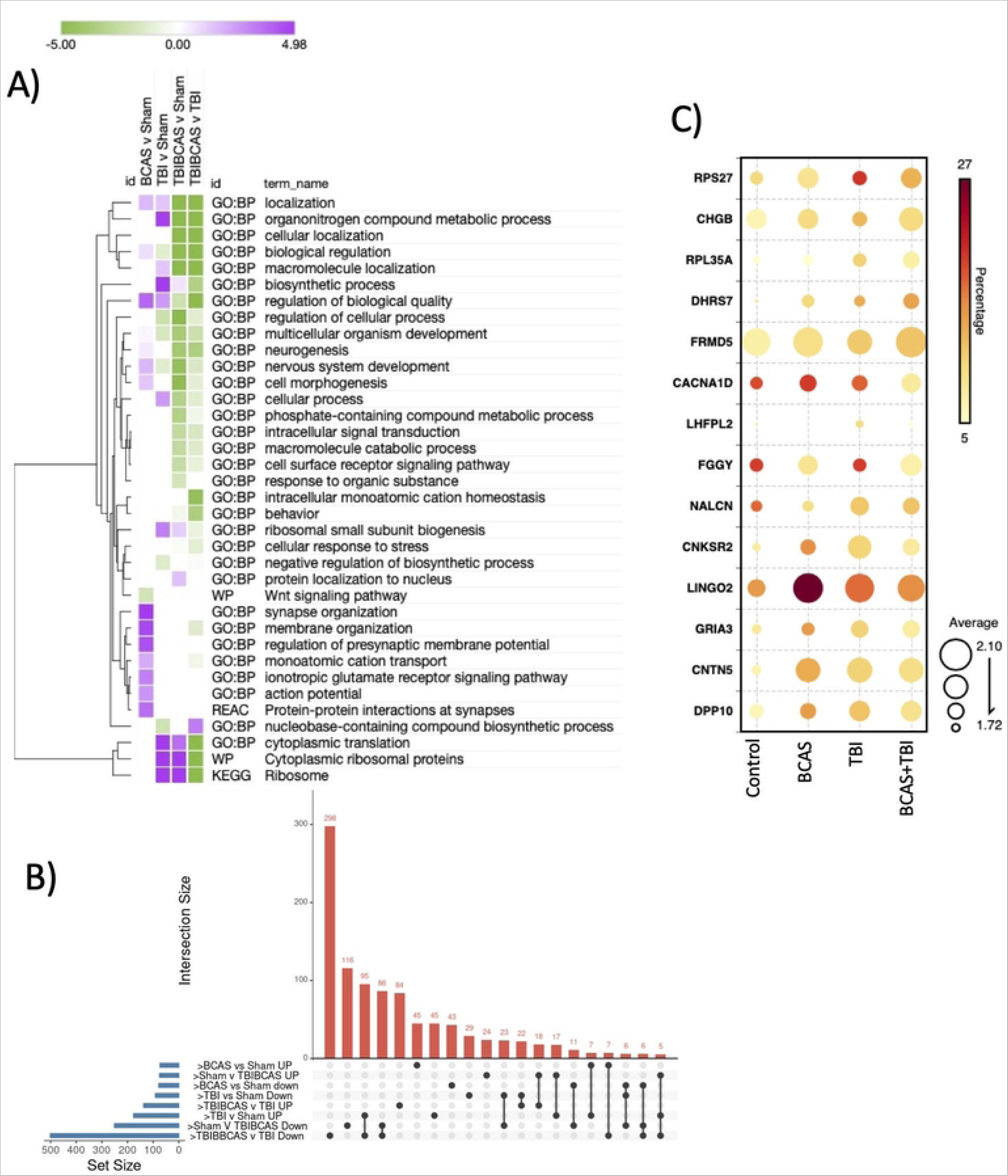
Transcriptomic signature in astrocytes. A) gene ontology terms derived from differentially expressed genes, B) upset plot and, C) notable gene changes.

#### Astrocytes

BCAS drove gene expression changes related to synaptic transmission and nervous system development in astrocytes (Fig 8). In contrast, TBI greatly upregulated ribosome-related genes and tended to reduce those related to cellular and nervous system development. BCAS+TBI significantly downregulated genes associated with cell surface signaling, neurogenesis, response to organic stimuli. While ribosome genes were upregulated relative to SHAM in BCAS+TBI, they were downregulated when comparing BCAS+TBI to TBI indicating that the ribosome gene induction was smaller in BCAS+TBI than TBI alone. BCAS+TBI also downregulated genes related to neurogenesis, nervous system development, and intracellular signal transduction.

**Fig 8.**
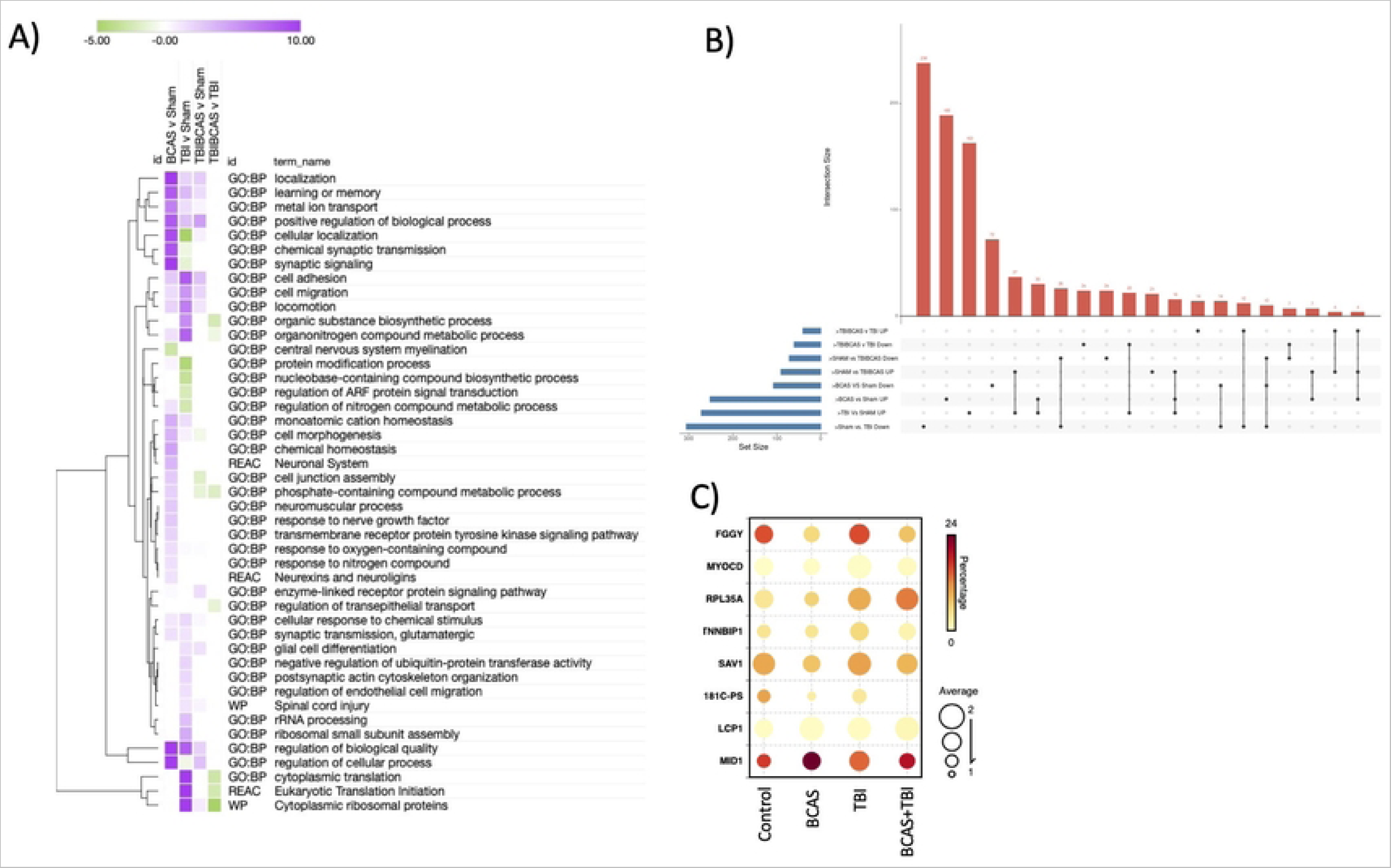
Transcriptomic signature in oligodendrocytes. A) gene ontology terms of DEGs, B) upset plot and, C) notable gene changes.

## DISCUSSION

There is growing evidence that traumatic brain injury represents the start of a disease process that interacts reciprocally with cardio- and cerebrovascular dysfunction. Here we tested the hypothesis that modest cerebral hypoperfusion would exacerbate the functional, vascular, histological, and transcriptomic responses to a mild-moderate TBI. Swiss-Webster mice of both sexes were implanted with steel microcoils around the carotid arteries bilaterally. The coils were .2mm in diameter, a size that has previously been shown to slightly and transiently impair cerebral blood flow(24). Thirty days later we induced a mild-moderate closed head traumatic brain injury. BCAS slightly reduced blood flow (around a 10% reduction relative to control mice) in both sexes at 30-days post implantation. TBI also caused a reduction in CBF, causing a reduction roughly similar in magnitude to BCAS. However, mice that underwent both procedures exhibited much larger reductions in CBF. This effect was paralleled by deficits in spatial learning and memory. Among males, BCAS+TBI animals exhibited the most severe deficits in Barnes maze performance. Surprisingly, among females there was an effect of BCAS on learning and memory, but not of TBI. Additionally, among males the combination of BCAS+TBI significantly exacerbated axonal damage relative to animals that experienced only TBI. Whereas among females no such additive effect of BCAS and TBI was apparent. Single nuclei RNA sequencing revealed that BCAS, TBI and the combination each produced unique, cell type specific adjustments in gene expression.

The mechanisms underlying reduced cerebral blood flow after traumatic brain injury remain poorly understood but may represent a homeostatic attempt to maintain tissue oxygenation. Indeed, oxygenation of brain tissue has long been conceptualized as determined by blood flow and capillary surface area and thus increases in cerebral blood flow and capillary diameter are directly proportional to tissue oxygenation (34, 35). However, this relationship does not explain the increase in oxygen extraction during functional activation which greatly exceeds increases in blood flow. Indeed, it ignores the biophysical reality that as cerebral blood flow increases, the time that oxygenated erythrocytes spend in capillaries decrease and the potential for oxygen extraction falls. Under resting conditions there is significant variability in the speed of blood flow through capillaries, termed capillary transit heterogeneity (CTH). Under healthy conditions capillary transit homogenizes as tissue metabolic needs increase (36). However, under conditions of vascular dysfunction, or other disease states where CTH cannot be decreased, cerebral blood flow can decrease (thus prolonging time red blood cells spend in capillaries) and paradoxically increase tissue oxygen extraction in order to meet tissue metabolic needs (37).

Acute alterations in cerebral blood flow are a common outcome of brain injury in both experimental animal studies and clinical populations (38–41). The general pattern is that blood flow declines acutely and then gradually recovers or in some cases overshoots pre-injury levels (42). Moreover, reduced cerebral blood flow is a strong predictor of poor TBI outcomes (43). In milder TBI cases, however, increases in oxygen extraction appears to compensate for the reduction in cerebral blood flow, thus providing sufficiently for tissue metabolic needs (44). The precise mechanisms for the acute reduction in blood flow after TBI remains poorly understood but is likely to result at least in part from edema, pericyte dysfunction and oxidant exposure. It also remains possible that the acute reduction in cerebral blood flow is necessary to maintain tissue oxygenation in the face of acute injury.

Here mice underwent BCAS, a manipulation that causes a relatively small reduction in cerebral blood flow. This procedure is very likely to increase CTH as both experimental animals and human carotid stenosis patients exhibit capillary dysfunction that is partially alleviated by improving perfusion (45, 46). Given the relatively mild reduction in CBF, it is likely that BCAS animals were able to maintain tissue oxygenation. Indeed, there was minimal evidence that the BCAS procedure produced gross neuropathology as evidenced by silver staining. Moreover, transcriptomic evidence of hypoxia or responses to reactive oxygen species were very limited, although since we sampled tissue 30 days after BCAS we cannot rule out the possibility of transient hypoxia-ischemia. TBI in BCAS mice persistently reduced CBF, an effect that was still apparent more than two weeks after injury. Thus BCAS+TBI animals maintained a level of blood flow lower than what can physically pass through the carotids. Whether this reduction in blood flow is a homeostatic response necessary for maintaining tissue oxygenation remains unspecified. However, what is known is that TBI is a metabolically expensive process that typically involves a period of increased metabolic substrate uptake. Limitations of these substrates greatly exacerbate disease outcomes. Whether the compromised vascular system of the BCAS animals can supply these metabolic fuels to the brains seems unlikely given the poorer functional and histological outcomes among BCAS+TBI animals relative to TBI.

Somewhat surprisingly females responded differently to the BCAS and TBI manipulations. Specifically, BCAS produced a milder reduction in cerebral blood flow. Sex differences have been reported in cerebral blood flow across the lifespan with females generally displaying greater CBF and sex steroid hormones are found throughout the vasculature (47). There have been relatively few studies that directly compare hypoperfusion outcomes across the sexes, however treatment with the sex steroid estradiol and progesterone both alleviate the deleterious consequences of hypoperfusion in male rodents (48, 49). Additionally, where there were clear detrimental effects on spatial learning and memory of both BCAS and TBI among males, females exhibited only a BCAS effect. The apparent sex difference in injury outcomes is not unexpected as females have been shown to be relatively protected from behavioral consequences of injury (50). Finally, there was no apparent increase in axonal degeneration among BCAS+TBI females relative to TBI. It remains unclear whether more severe brain injuries would be exacerbated by BCAS among females.

Given the prominent interactions between BCAS and TBI among males we elected to perform single nuclei RNA sequencing on the males only. The overall patterns of gene expression varied markedly between cell types and experimental conditions. For instance, many of the genes altered by BCAS alone, including those for cellular development and plasticity, were not changed in the BCAS+TBI group. Among both excitatory and inhibitory neurons for instance, BCAS+TBI produced dramatic upregulation of genes related to ribosome biogenesis and oxidative phosphorylation relative to SHAM. However, what is surprising is that these genes associated with protein generation and metabolic activity were not induced by TBI alone. Indeed, among GABAergic neurons TBI reduced the expression of genes related to the electron transport chain and oxidative phosphorylation. Thus, it seems possible that neurons, at least at this timepoint, are not directly responsible for the elevated metabolic activity often reported after TBI.

Glial cell responses to BCAS and TBI were different than neurons. BCAS astrocytes upregulated a number of genes related to synaptic function, plasticity, and development. These synaptic genes were not upregulated in BCAS+TBI astrocytes (or were downregulated in some cases) indicating that synaptic modulations induced by BCAS were reversed in animals that were subsequently injured. Moreover, ribosomal subunit gene expression was upregulated by TBI and BCAS+TBI (although the upregulation was of a greater magnitude among TBI only mice as indicated by a downregulation of ribosome-related genes when comparing TBI and BCAS+TBI). Also, in contrast to neurons, the largest overall effects of BCAS+TBI among astrocytes was a suppression of gene expression and consequent downregulation of genes relating to signaling, plasticity, and biological regulation. Many genes were also upregulated in oligodendrocytes in response to BCAS, including those associated with synaptic plasticity and other homeostatic processes which were lost in BCAS+TBI. Oligodendrocytes upregulated ribosome genes following TBI, but this effect was largely absent among BCAS+TBI animals.

It should be pointed out that inflammatory responses are notably absent from gene expression profiles in cell types where we might expect them (e.g., astrocytes and microglia). Indeed, we recovered relatively few microglia from our isolation and the gene expression profiles were not altered significantly by our manipulations (data in supplement). Still, it remains puzzling that this set of manipulations produced little in the way of inflammatory signaling, at least at this time point.

Thus, the patterns of transcriptional activity varied in a cell type- and condition dependent manner. One illustrative example is ribosome subunit gene expression, this category of gene expression is typically associated with cellular activation and preparation for protein production, both of which are metabolically expensive processes. Among neurons, only BCAS+TBI animals exhibited an upregulation of these proteins. In contrast, astrocytes exhibited strong upregulation of ribosome-related genes following TBI regardless of BCAS while oligodendrocytes only upregulated ribosome genes following TBI in the absence of BCAS. Upregulation of ribosome related genes has been reported in blast and fluid percussion injuries previously and appears at least partially to be sensitive to depletion of microglia with CSF1R inhibitors (51–53). What is somewhat surprising however, is that cells that are presumably in a resource-challenged environment after BCAS+TBI would engage in an energetically expensive process that does not seem to occur in animals that only underwent TBI. Moreover, excitatory neurons in the BCAS+TBI group also upregulated genes related to aerobic respiration suggesting that metabolic needs of the tissue were not being met. Gene expression changes are unlikely to represent the TBI being insufficient to activate damage-related programs as there were significant gene expression shifts following TBI in glial cells and functional deficits and axonal damage even in the absence of BCAS. One significant limitation of this approach, however, is that RNA sequencing was conducted at a single time point, we are thus unable to determine whether there are different temporal patterns of gene expression in different cell types and perhaps cells from TBI only groups undergo similar adjustments but were not detected.

Taken together these gene expression changes provide some overall clues to the events that culminate in poor spatial learning and memory and axonal degeneration after BCAS+TBI. Specifically, at the time point we measured 30 days after BCAS implantation there were marked shifts in gene expression among virtually all cell types, many of which appear to be homeostatic responses to the adjustments in blood flow. Indeed, this mild BCAS procedure appeared to be well tolerated, at least by males, as we detected no deleterious consequences for spatial learning and memory or axon damage. Moreover, at this time point 24 hours after injury there were surprisingly few gene expression shifts induced by TBI alone. However, when these manipulations were combined and mice with BCAS coils were injured, the situation was quite different. Many of the gene expression shifts that appeared compensatory to challenge were lost and cells engaged in overall patterns of gene expression that were notably different from either BCAS or TBI alone. Whether the reduction in cerebral blood flow *per se* or the adjustments that cells made to respond to this reduction was responsible for these consequences remains unspecified.

There are several individual genes that merit specific mention and future attention. Myocardin (MYOCD) is a transcription factor that was originally described in cardiac muscle cells and is involved in serum response factor (SRF). Moreover, activation of the SRF-MYOCD axis in brain vascular smooth muscle cells results in arterial hypercontractility, cerebral hypoperfusion and the accumulation of A-beta vessels. We detected MYOCD specifically among BCAS+TBI animals in neurons (54). However, there is little extant evidence of MYOCD being expressed outside of muscle cells and thus the role of this gene here remains unspecified. Midline1 (MID1) is a gene that was originally discovered as the cause of Opitz-BBB/G syndrome (OS) a disease with multiple malformations of the ventral midline. MID1 gene expression was strongly upregulated by BCAS in endothelial cells and oligodendrocytes. The protein encoded by MID1 is a member of the RING finger family and has a multitude of functional domains including, an E3 ubiquitin ligase and a variety of others responsible for protein-protein and protein-nucleic acid binding (55). MID1 has more recently been linked to neurodegeneration and inhibition of MID1 activity reduces amyloid precursor protein expression in cultured primary neurons (56). FGGY encodes a carbohydrate kinase which is induced by TBI across neurons and glial cells and was previously identified as a marker gene for neurodegeneration in a genome-wide association study (57).

In conclusion, the study underscores the complexity of interactions between cerebral hypoperfusion and TBI. While each condition individually leads to specific vascular, functional, histological, and transcriptomic changes, their combination results in outcomes that aren’t simply additive, pointing towards intricate interplay at molecular, cellular, and system levels. This work also emphasizes the importance of considering sex as a biological variable, as males and females exhibited different responses to the experimental manipulations. Further research is required to delve deeper into the mechanistic insights and the temporal dynamics of these observed changes, which could potentially aid in developing therapeutic interventions for TBI and cerebrovascular dysfunctions.

## Acknowledgments

This work was funded by the WV Stroke CoBRE grant (5P20GM109098-08) and the WVU Center for Foundational Neuroscience Research and Education pilot award. The authors thank Dr. A. Courtney DeVries for her ongoing advice and support.

## Data availability

The snRNA sequencing data generated in the current study are available under GEO accession number GSE241690. Data are available for reviewers at https://www.ncbi.nlm.nih.gov/geo/query/acc.cgi?acc=GSE241690 with token cjmzkcoyznkdvif.

